# Rewiring of an ancestral Tbx1/10-Ebf-Mrf network for pharyngeal muscle specification in distinct embryonic lineages

**DOI:** 10.1101/039289

**Authors:** Theadora Tolkin, Lionel Christiaen

**Author notes:** **Summary Statement:** We adapted the heterologous LexA/LexAop system in Ciona, and used it to characterize rewired myogenic programs in separate embryonic lineages.

## Abstract

Skeletal muscles arise from diverse embryonic origins, yet converge on common regulatory programs involving muscle regulatory factor (MRF)-family genes. Here, we compare the molecular basis of myogenesis in two separate muscle groups in the simple chordate *Ciona:* the <u>a</u>trial and <u>o</u>ral <u>s</u>iphon <u>m</u>uscles. Here, we describe the ontogeny of OSM progenitors and characterize the clonal origins of OSM founders to compare mechanisms of OSM specification to what has been established for ASM. We determined that, as is the case in the ASM, *Ebf and Tbx1/10* are both expressed and function upstream of *Mrf* in the OSM founder cells. However, regulatory relationships between *Tbx1/10, Ebf* and *Mrf* differ between the OSM and ASM lineages: while *Tbx1/10, Ebf* and *Mrf* form a linear cascade in the ASM, *Ebf* and *Tbx1/10* are expressed in the inverse temporal order and are required together in order to activate *Mrf* in the OSM founder cells.

## Introduction

In Vertebrates, differentiation of skeletal muscle during development relies on the deployment of a core myogenic network comprising the MRF-family of bHLH transcription factors, *MyoD, Myf5*, and *Mrf4* (Block and Miller 1992; Rudnicki et al. 1993; Braun et al. 1994; Summerbell, Halai, and Rigby 2002; Kassar-Duchossoy et al. 2004). However, the exact relationships between these myogenic genes in activating expression of the muscle-specific differentiation program, as well as the factors that specify myogenic potential upstream of the core MRF-based network, differ between regions of the embryo (Sambasivan et al. 2009; Harel et al. 2009; Adachi et al.; Buckingham & Rigby 2014). For example, while *Pax3* is an important muscle determinant in somitic lineages, *Tbx1* determines branchiomeric muscle, and *Pitx2* is an activator of the core myogenic network in extraocular muscles (Harel et al. 2009; Sambasivan et al. 2009).

Research in *Drosophila, C. elegans*, sea urchins, Amphioxus, and non-model systems including Spiralians, Cnidarians, and brachiopods has shown that MRF-family bHLH transcription factors have roles in muscle development across Deuterostomes, and even throughout Bilaterians (Andrikou et al. 2015; Buckingham & Rigby 2014; Schaub et al. 2012;Ciglar & Furlong 2009; Zhang et al. 1999). Moreover, there is evidence, discussed below, that upstream regulatory factors responsible for *MRF* expression and muscle development may be conserved across long evolutionary distances. In particular, the COE (Collier/Olfactory-I/Early B-Cell Factor) family of transcription factors has been implicated in striated muscle development in *Drosophila* (Crozatier and Vincent 1999), *Ciona* (Stolfi et al., 2010; Razy-Krajka et al. 2014), *Xenopus* (Green and Vetter 2011) and chick (El-Magd et al., 2013, 2014a, 2014b). Homologs of the T-box transcripton factor Tbx1 also seems to have a role in muscle identity in *Drosophila* (Schaub et al. 2012), *Ciona* (Wang et al. 2013), mice (Aggarwal et al. 2010) and humans (H. Yagi et al. 2003).

In the basal chordate *Ciona*, distinct muscle groups also arise from different blastomeres in the early embryo, and the single MRF-family gene in the *Ciona* genome, called *Mrf*, is expressed in all non-cardiac muscle precursors (Meedel, Chang, and Yasuo 2007; Razy-Krajka et al. 2014). In the B7.5 lineage, *Mrf* is first transiently expressed in B7.5 and in the B8.9 and B8.10 founder cells, before induction of the cardiopharyngeal progenitors (called trunk ventral cells, TVCs) (Christiaen et al. 2008). *Mrf* is later reactivated downstream of *Ebf* specifically in the atrial siphon muscle (ASM) founder cells. ASMFs give rise to *Mrf+* differentiating muscle cells as well as *Bhlh-tun1+/Mrf*-stem-like muscle precursors, which cease to express *Mrf* in response to Notch-mediated lateral inhibition. The progeny of these *Bhlh-tun1*-expressing cells reactivate *Mrf* after metamorphosis in order to give rise to the body wall muscles of the juvenile (Razy-Krajka et al. 2014).

The oral siphon muscles (OSM) derive from a distinct lineage, named A7.6, which is not cardiogenic but instead also produces blood and tunic cells, as well as stomach and gill slit epithelia (Hirano and Nishida, 1997; Tokuoka et al. 2005). Previous *in situ* analyses of ASM-speciflc gene expression indicated that most ASM-expressed genes, including the key regulators *Ebf* and *Mrf*, were also active in the OSM (Razy-Krajka et al., 2014). We sought to characterize the lineage of A7.6-derived OSM precursors, as well as regulatory mechanisms leading to the activation of a core siphon muscle program in the OSMs. Because no A7.6-specific reporter transgene was available, we applied the heterologous, repressible transgenic systems Gal4/UAS (Brand and Perrimon 1993), LexA/LexAop (R. Yagi, Mayer, and Basler 2010), TrpR/tUAS (Suli et al. 2014), and QF/QS (Potter et al. 2010) and systematically evaluated their efficacy, specificity and toxicity for use by multiplexed electroporation in *Ciona* embryos. We identified a Gal80-repressible form of LexA/LexAop (R. Yagi, Mayer, and Basler 2010) to be the most efficient, specific and innocuous approach to specifically mark the A7.6 lineage.

Using a defined combination of LexA, LexAop and Gal80 transgenes, we report a detailed description of the ontogeny of the A7.6 lineage, the progeny of which has previously been referred to as the trunk lateral cells (TLC; Tokuoka et al. 2004; Imai et al. 2003; Tokuoka et al. 2005; Satou et al. 2001). We identified each stereotypical division of the anterior-most descendants of the TLC, leading to the birth of two sister, fate-restricted, OSM founder cells (OSMF), on either side of the embryo. We find that OSM commitment is defined by expression of *Ebf* and *Tbx1/10* exclusively in the OSMFs, which both activate *Mrf* but produce a mixed population of OSM precursors (OSMPs) expressing either *Mrf* or *Bhlh-tun1*, as is the case in differentiating or stem-cell-like ASMPs, respectively. Next we demonstrate that, in contrast to what has been shown in the ASM, *Mrf* expression in the OSM rudiment depends on the joint activities of *Ebf* and *Tbx1/10*. Thus, while *Tbx1/10* regulates *Ebf* in the ASMF where *Ebf* acts as a master-regulator of ASM fate, *Ebf* is regulated independently of *Tbx1/10*, and both are required jointly for *Mrf* expression in the OSM. Our findings demonstrate context-dependent rewiring of a deeply conserved kernel of muscle specification genes within a single genome.

## Results

### Heterologous binary systems for lineage-specific transgene expression in Ciona

Many genetic studies in *Ciona* are based on the use of transient transgenesis by electroporation of plasmid DNA (Corbo, Levine, and Zeller 1997; Christiaen et al. 2009a; Stolfi and Christiaen 2012). Putative enhancers are cloned upstream of neutral reporters or active genetic elements for tissue-specific transgene expression, generally recapitulating endogenous patterns of activity (Wang and Christiaen 2012). Pleiotropic *cis*-regulatory regions can be dissected to isolate tissue-specific enhancers, but this process is not guaranteed to generate a transgene with specific enough enhancer activity.

Heterologous transgene activation systems provide an attractive addition to the molecular toolbox for studies using *Ciona*. Gal4/UAS-based systems can be inhibited by expression of Gal80, generating a logic by which tissue specificity is achieved by activation of the response element within the domain of gene A, “but not” within the domain of partially overlapping gene B (Brand and Perrimon 1993; Suster et al. 2004).

We chose the proximal enhancer of *Hand-related*, which is expressed in A7.6 and anterior trunk endoderm (Davidson and Levine 2003; Woznica et al 2012; and this paper) to test the specificity, efficiency, and toxicity of the heterologous, repressible transgenic systems Gal4/UAS (Brand and Perrimon 1993), LexA/LexAop (R. Yagi, Mayer, and Basler 2010; Lai and Lee 2006), TrpR/tUAS (Suli et al. 2014), and QS/QF (Potter et al. 2010) (Fig. S1). Because of its rapid activation, efficiency, specificity, lack of obvious toxicity, and ability to be inhibited by Gal80, we used the Gal80-repressible form of a transcription activator consisting of the DNA binding domain from the bacterial LexA protein fused to the trans-activation domain of the yeast Gal4 protein, LexA::Gal4AD (“LHG”) (R. Yagi, Mayer, and Basier 2010) downstream of the *Hand-related* proximal enhancer (Hand-r>LHG). We electroporated zygotes using a combination of *Hand-r>H2B::mCherry;Hand-r>LHG;LexAop>GFP* constructs in order to determine the extent to which the binary LexA/LexAop system recapitulated the original *Hand-r*-driven expression pattern (Fig. 1A,B). We observed that 83.1% (± 17.3%; n=107) of larvae expressing *Hand-r>H2B::mCherry* also expressed *Hand-r>LHG;LexAop>GFP* in patterns that overlapped almost completely, meaning that the LexA/LexAop binary system faithfully recapitulates the activity of unary transgenes, albeit with a ∼3 hours delay compared to reach levels similar to those of unary *Hand-r*-driven transgenes (Fig. S1C).

**Figure 1.**
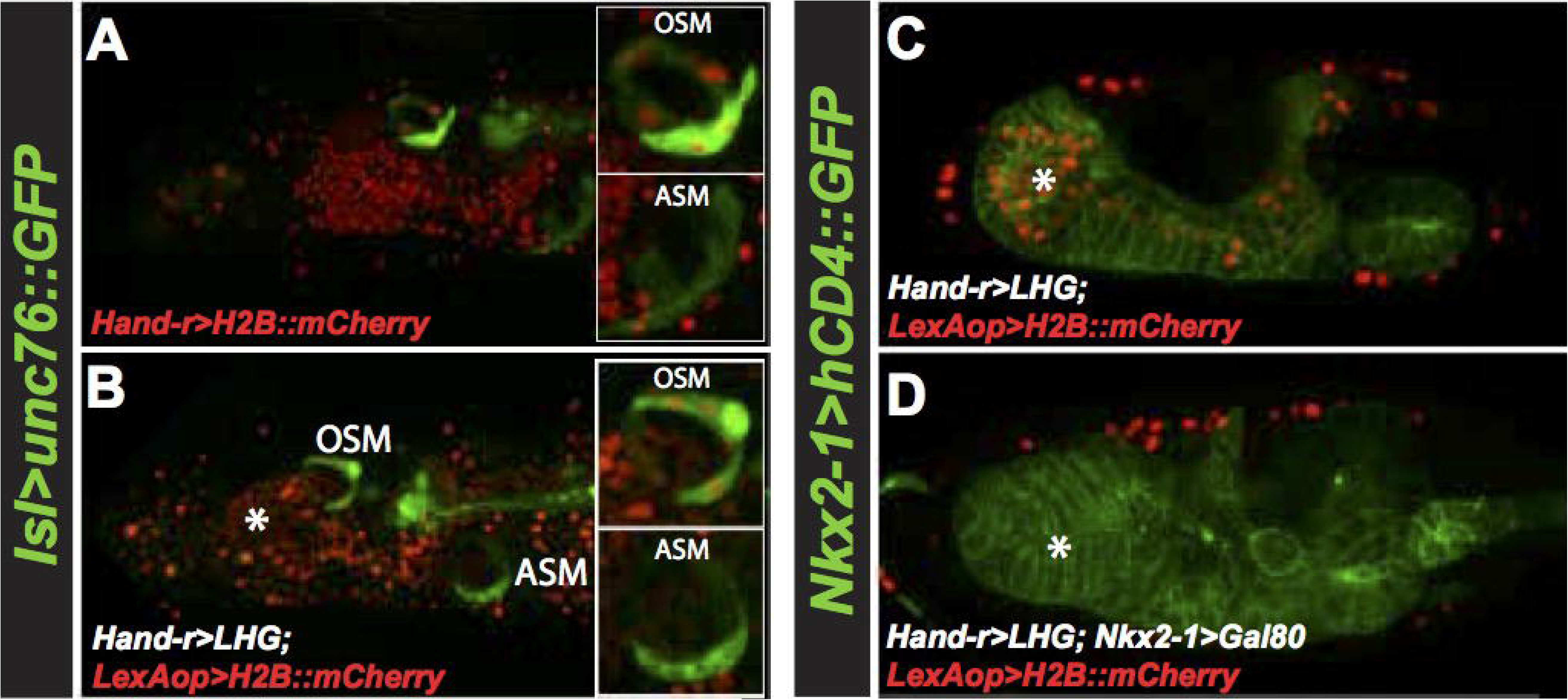
The Gal80-repressible LexA/LexAop binary transgenic system efficiently and specifically labels the A7.6 blastomeric lineage of *Ciona robusta*. (A) Confocal image of 24hpf larva showing expression of Handr>H2B:mCh and Isl>unc76:GFP. OSM are mCherry+/GFP+ while ASM are mCherry-/GFP+. Endoderm is marked with “*”. (B) 24hpf larva expressing Hand-r>LHG;LexAop>H2B:mCherry and Isl>unc76:GFP, showing that Handr>LHG;LexAop>unc76:GFP fully recapitulates expression of Handr>H2B:mCherry. (C) Single confocal slice of 24hpf larva expressing Hand-r>LHG;LexAop>H2B:MCherry and the endodermal marker Nkx2-1>CD4:GFP, showing that Hand-r>LHG;LexAop>H2B:mCherry is expressed throughout the endoderm. (D) Single confocal slice of 24hpf larva showing that expression of Nkx2-1>Gal80 abolishes expression of Hand-r>LHG;LexAop>H2B:mCherry in the endoderm only.

We then used the *Nkx2-1/Ttf1* enhancer to drive Gal80 (*Nkx2-1>Gal80*) in order to inhibit LHG trans-activity specifically in the endoderm and restrict *LexAop* activation to the A7.6 lineage (Figure 1C, D; Ristoratore et al., 1999). We marked the endoderm membranes with *Nkx2-1>hCD4::GFP* (Gline et al. 2015), and used either *Hand-r>LHG;LexAop>H2B::mCherry* or *Hand-r>LHG;Nkx2-1>Gal80;LexAop>H2B::mCherry*. We observed a 96% reduction in the proportion of hCD4::GFP+ larvae that also had endodermal cells marked with H2B::mCherry (Fig. 1C,D). These results demonstrate the efficacy and specificity of the LHG/Gal80/LexAop system to achieve refined transgene expression using a tertiary “but not” logic. Therefore, the combination of *Hand-r>LHG; Nkx2-1>Gal80;LexAop>GFP* constructs (hereafter referred to as LexO(A7.6)≫GFP for simplicity) drives transgene expression specifically in the A7.6 lineage and its descendants.

### The detailed A7.6 origins of oral siphon muscles (OSM)

Although previous work has characterized gene expression patterns attributed to the A7.6-derived TLCs in tailbud embryos (Imai, Satoh, and Satou 2003; Jeffery et al. 2008; Shi and Levine 2008; Tokuoka et al. 2004; Tokuoka, Satoh, and Satou 2005), the detailed development and morphogenesis of the A7.6-derived cells remained uncharacterized. Therefore, we undertook a comprehensive description of the entire A7.6 lineage through 8.5 hours post-fertilization (hpf; stage 17, initial tailbud I;(Hotta et al. 2007)), and of the cells that give rise to the OSM through 24hpf (stage 26+).

Because mCherry proteins generated by the LexA/LexAop constructs are not easily detectable in embryos until approximately 8hpf (Fig. S1C), we marked the A7.6 lineage by the overlap of *Hand-r>H2B:mCherry* and *MyT>unc76:GFP* transgenes, (The *MyT* driver is active in A7.6 and a-line neural cells (Imai et al. 2004; Shi and Levine 2008); Fig. 2B-F). We visualized the A7.6 blastomeres in samples fixed every fifteen minutes starting at 5hpf (approx. stage 10), and applied Conklin's nomenclature (Conklin 1905) to identify and name every cell within the A7.6 lineage up to 8.5hpf (approx. stage 17; Fig. 2). In parallel, we tested whether early divisions coincided with specific gene activities, as is often the case in early ascidian embryos (Satoh 2013, ch. 12): we performed *in situ* hybridization on early stage embryos to detect expression of *Hand-related* and *MyT* immediately following each one of the earliest divisions, while marking A7.6-derived cells with *MyT*-driven nuclear H2B::mCherry *(MyT>H2B::mCherry)* (Fig. 2G-P).

**Figure 2.**
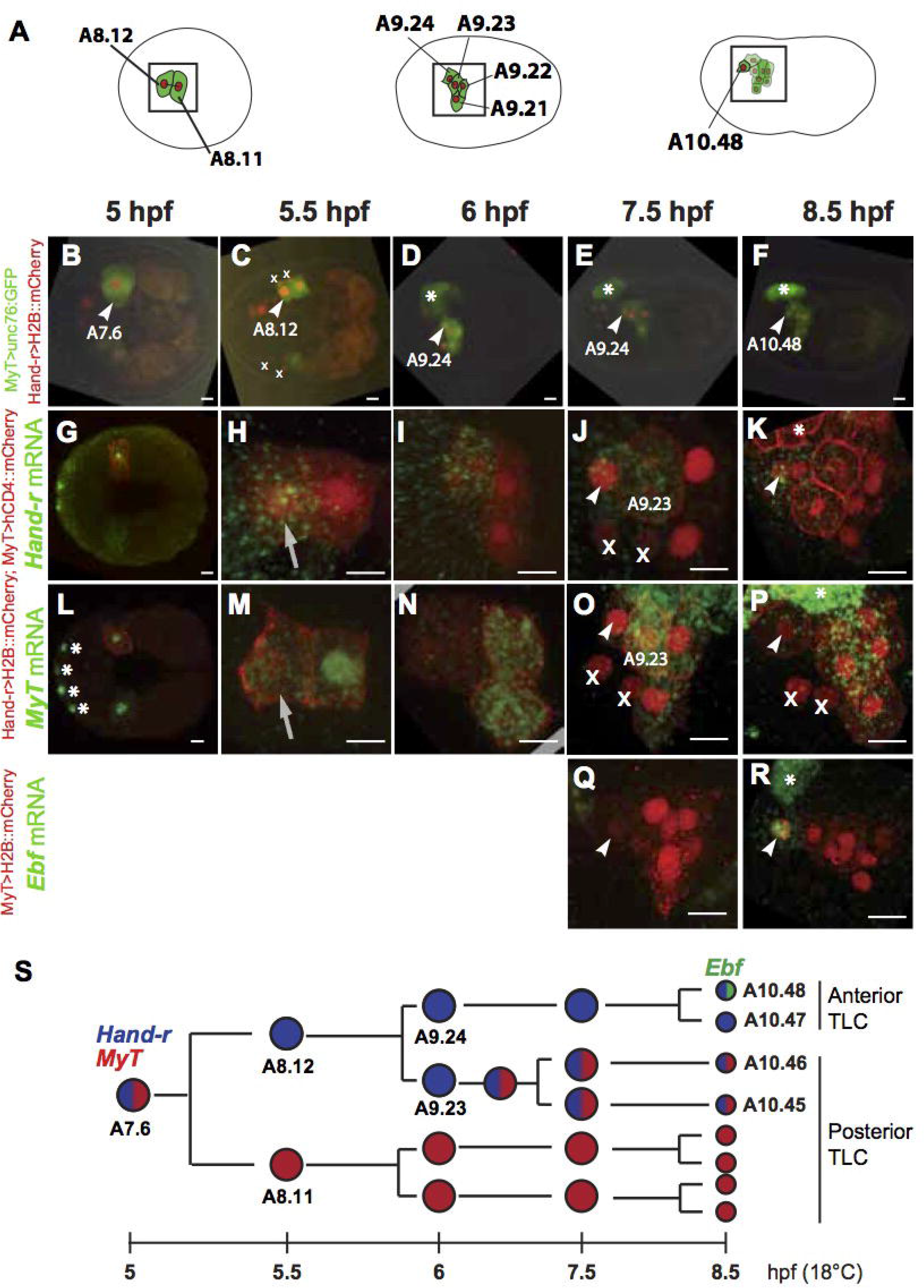
Embryonic development of the trunk lateral cells (TLC) 5.5hpf to 8.5hpf. (A) Cartoon of divisions within the A7.6 lineage at 5.5hpf, 6hpf and 8.5hpf. Each cell is named according to the scheme developed by Conklin (Conklin 1905). Boxes indicate the regions shown at each time-point in panels G-R. (B-F) Embryos electroporated with Hand-r>H2B:mCherry and MyT>unc76:YFP, which overlap exclusively in the A7.6 lineage. Note the faint expression of H2B:mCherry in two adjacent endoderm cells beginning at 5.5hpf, presumably derived from A7.5, the sister of A7.6, thus reflecting an occasional early onset of transgene expression in the mother A6.3. These A6.3-derived cells are marked with a Roman numeral “x” wherever they appear. This staining was accounted for in all observations, and did not interfere with our ability to identify A7.6-derived cells. Expression of MyT>GFP or of *MyT* mRNA in the a-line neural cells is marked with an “*”. (GR) Expression dynamics of *Hand-related* (G-K), *MyT* (L-P), or *Ebf* (Q,R) mRNA, with A7.6 lineage marked by MyT>H2B:mCherry. (S) Schematic diagram of all TLC divisions from 5hpf to 8.5hpf, with gene expression patterns mapped on to each cell and color-coded. Blue = *hand-related;* Red = *MyT;* Green = *Ebf*. Scale bars=25μm.

At 5hpf, newborn A7.6 blastomeres express both *Hand-r* and *MyT* (Fig. 2G,L); and by 5.5hpf, they have divided along the antero-posterior axis (Fig. 2C, H, M). Although there is no obvious morphological asymmetry between the two daughter cells, the anterior A8.12 preferentially maintains *Hand-r* expression, while the posterior A8.11 maintains *MyT* expression (Fig 2H,M). By 6hpf, both A8.12 and A8.11 have divided; A8.12 along the anteroposterior axis and A8.11 dorsoventrally, again without obvious morphological asymmetry between their daughter cells, A9.21/A9.22 and A9.23/A9.24, respectively (Fig. 2A,D). A9.24/A9.23 both maintained roughly equal levels of *Hand-r* expression, which was no longer detected in the posterior A9.22/A9.21pair, which instead expressed sustained levels of MyT (Fig. 2I,N).

*MyT* expression was rapidly and strongly re-activated in A9.23, the sister of the anteriormost *Hand-r+* A9.24 (Fig. 20), prior to dividing at ∼7.5hpf. Therefore, A9.23 expresses both *MyT* and *Hand-r* when it divides at ∼7.5hpf. This division occurs along the dorsoventral axis of the embryo, giving rise to the ventral A10.45 and dorsal A10.46 cells, both of which maintained *Hand-r* and *MyT* expressions until about 8.5hpf, at which point expression of *Hand-r* was decreased substantially (arrows in 2H and M point to A8.12, which maintains *Hand-r*, but not *MyT)*. By 8.5 hpf, the anteriormost TLC, A9.24, had divided along the anteroposterior axis and its daughters A10.48/47 maintained expression of *Hand-r* (Fig. 2K). At that time, which is during the late neurula stage, the anteriormost TLC, A10.48 activates *Ebf* (Fig. 2K,P,R). In summary, A7.6 blastomeres and their anterior-most progeny always divided somewhat antero-posteriorly such that A10.48, the anterior-most *Hand-r+/Ebf+* great-granddaughter of A7.6, always stood out among the TLCs. Given the established roles of *Hand-r* and *Ebf* in ASM specification within the cardiogenic B7.5 lineage (Stolfi et al., 2010; Razy-Krajka et al., 2014), we regarded A10.48 as the most likely OSM progenitor in tailbud embryos.

Between 8.5hpf and 16hpf, although the derivatives of A9.23 and A9.22/21 can still be distinguished from each other by position and gene expression patterns, more detailed clonal relationships could not be inferred from time series of fixed embryos. Therefore, we refer to all A9.23/22/21 derivatives as “posterior TLC”. By contrast, the A9.24 derivatives, A10.48 and A10.47, which we will call the “anterior TLC”, continue to follow stereotyped division patterns and gene expression dynamics until 16hpf (stage 24; LTBII; Fig. 3).

**Figure 3.**
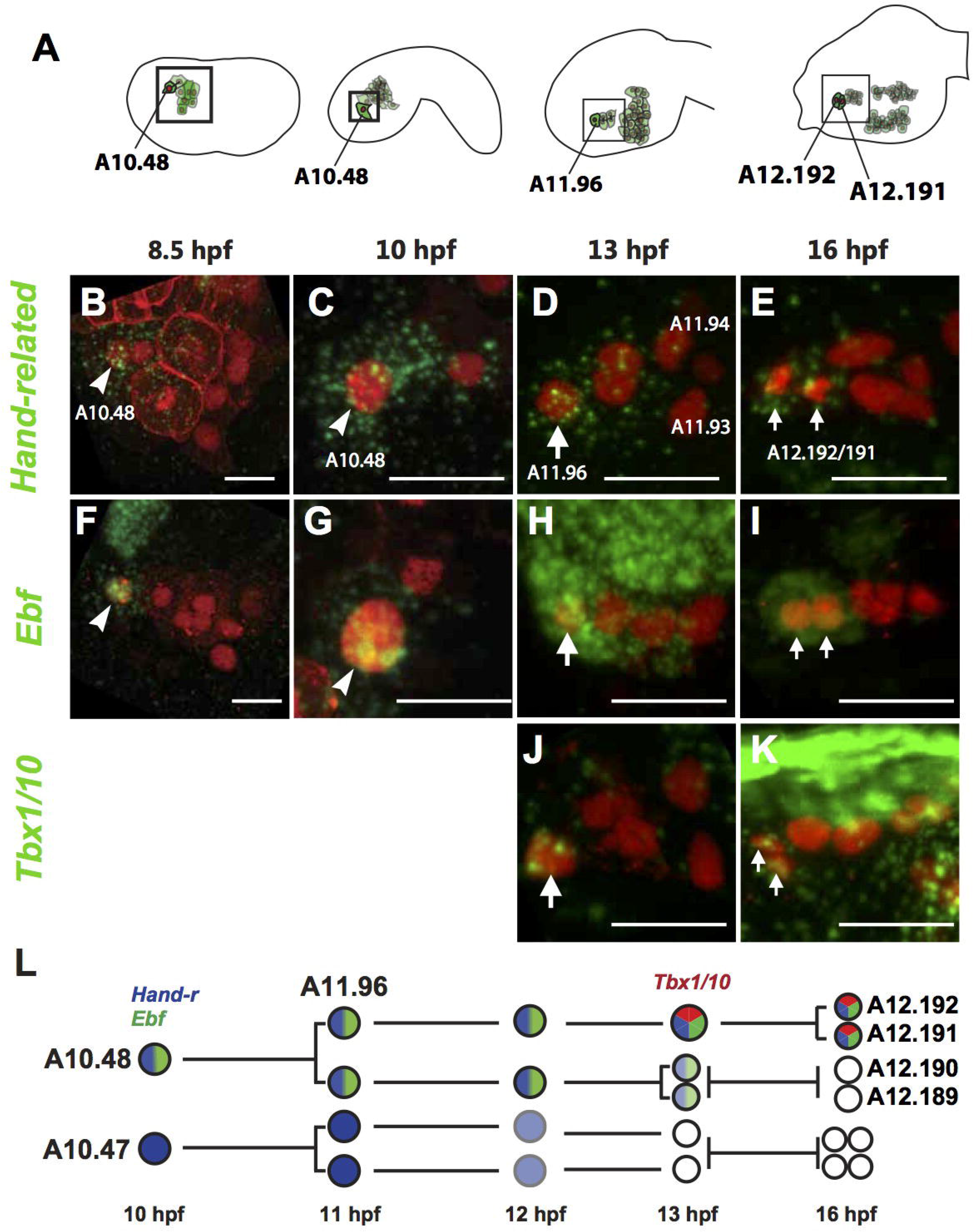
Embryonic development of the trunk lateral cells (TLC) 8.5hpf to 16hpf. First cartoon in A, and panels B and F are the same data shown in Fig. 2K and R, to emphasize continuity of A10.48 cell and to show the anterior TLC in the whole-embryo context. Panels B-K show only the anterior TLC, which are boxed in the cartoons in A. (B-K) Close-up of anterior TLC (descendants of A10.48/47 cell pair only) marked with LexO(A7.6)>>H2B:mCherry showing expression of *Hand-r* (B-E), *Ebf* (F-I) and *Tbx1/10* (J, K) at 8.5hpf, lOhpf, 13hpf, and 16hpf. Arrowheads mark the A10.48 cell; large white arrows mark A11.96; small white arrows mark the daughters of A11.96, named A12.192/191. Scale bars = 25 μm. (E) Schematic diagram of cell divisions and gene expression in the anterior TLC 10-16hpf. Blue = *hand-related;* Green = *Ebf;* Red = *Tbx1/10;* lighter versions of each color indicates lower levels of gene expression.

In order to test whether the OSM derive from the anterior *Hand-r+/Ebf+* A10.48 progenitors, we sought to further characterize the division, migration and gene expression patterns of the latter throughout tailbud and larval stages (Figs. 3, 4). For time-points after 9hpf, we used our *LexO(A7.6)≫H2B::mCherry* binary reporter system to label the A7.6 lineage, and applied the Conklin nomenclature only to the A10.48 cells and their descendants. Finally, It has been shown that *Hand-r, Tbx1/10* and *Ebf* are expressed sequentially and are necessary to activate the ASM program in B7.5-derived progenitors (Wang et al. 2013; Razy-Krajka et al. 2014; Stolfi et al. 2014). Therefore, we added *Tbx1/10* to *Hand-r* and *Ebf* in our analysis of candidate markers of early fate-restricted OSM precursors.

**Figure 4.**
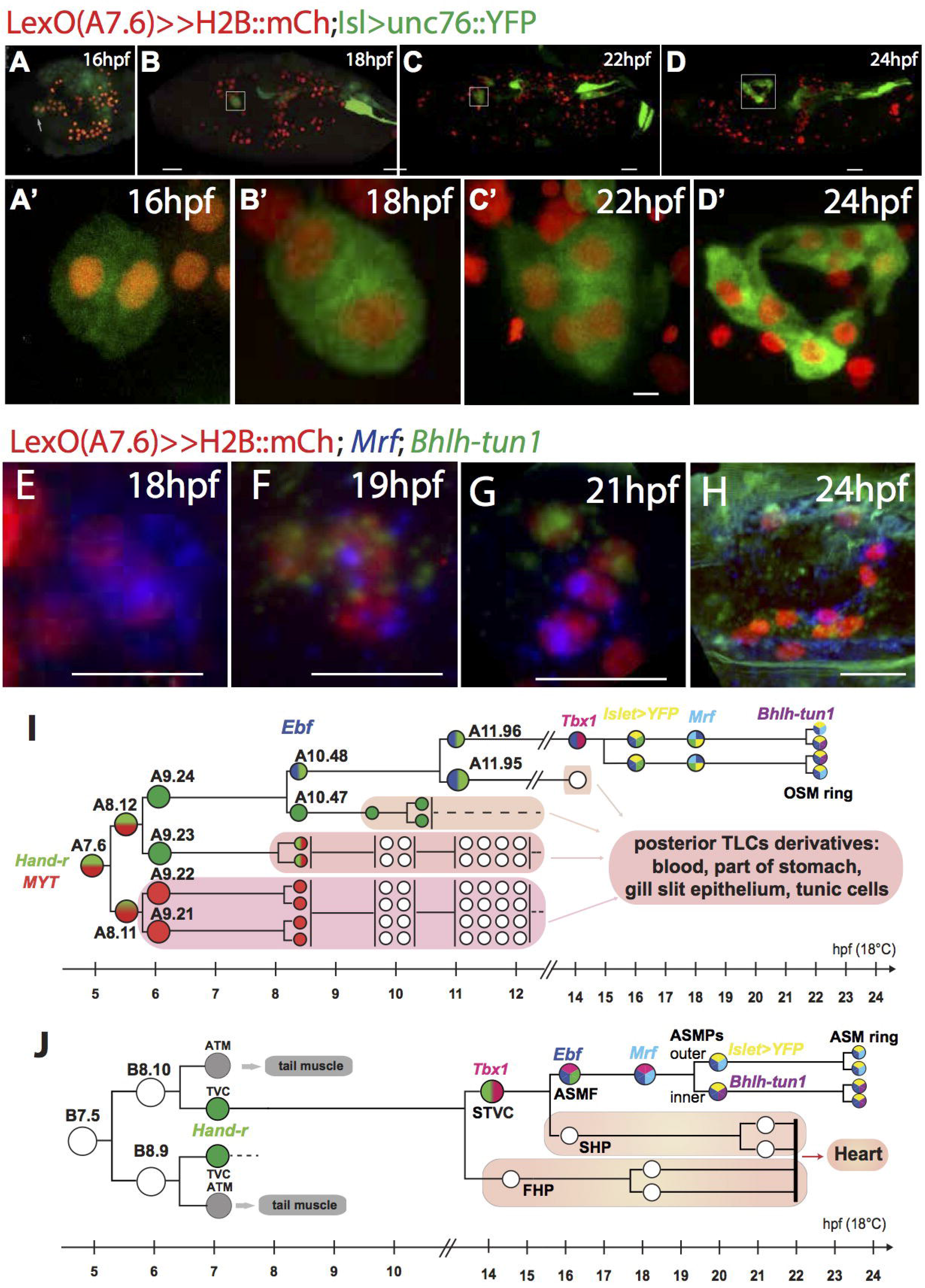
Development of the OSM precursors after larval hatching. (A-D) Larvae electroporated with LexO(A7.6)>>H2B:mCherry;Isl>unc76:GFP showing initiation of Isl>unc76:GFP expression in the OSMP at 16hpf (A), followed by cell divisions, anterior migration of the OSMP, and ring formation around the stomodeum by 24hpf (B-D). Boxed regions in A-D are shown close-up in A’-D’. (E-H) Close-up of OSMP in 18-24hpf larvae labeled with LexO(A7.6)>>H2B:mCh and with *Mrf* (blue) and *Orphan-bHLH-1* (green) mRNA revealed by *in situ* hybridization. (I) Complete schematic diagram of the development of the A7.6 lineage, from the time of its birth at 5hpf, until differentiation of the OSM around 24hpf, with gene expression dynamics labeled and mapped on clonally. (J) For comparison, a simplified schematic diagram of key steps in ASM development within the B7.5 lineage based on (Davidson and Levine 2003; Stolfi *et al*. 2010; Wang *et al*. 2013).

The *Hand-r+/Ebf+* A10.48 did not divide until 11hpf (mid-tailbud II), at which point its daughter cells, the anterior A11.96 and posterior A11.95, exhibit distinct and stereotyped differences in nuclear volumes and positions. Live imaging showed that this size asymmetry resulted from an inflation of A11.95 during interphase rather than an initially asymmetric division (movie S1). Meanwhile, A10.47 divides dorsoventrally giving rise to A11.94/93. During this time, both A11.96 and A11.95 continue to express *Hand-r* and *Ebf* while A11.94/93 lost the *Hand-r* mRNAs that were still detectable in their A10.47 mother (Fig. 2K; Fig. 3B-E,L). By 13hpf, A11.95 has divided dorsoventrally and both daughters A12.190 and A12.189 continue to express *Hand-r* and *Ebf* so that all three of A11.96, and A12.190/A12.189 co-express *Hand-r* and *Ebf*. However, A11.96 expresses higher levels of *Ebf* mRNAs than the A12.190/189 pair (Fig. 3H). Moreover, by 13hpf, only A11.96 activates *Tbx1/10* expression (Fig. 3J). By 16hpf, A11.96 has divided along the anteroposterior axis, giving rise to anterior A12.192 and posterior A12.191. The remaining anterior TLC have also divided, such that there were always 8 anterior TLCs by 16hpf, with only the anterior-most A12.192 and A12.191 cells co-expressing *Hand-r, Ebf and Tbx1/10* (Fig. 3E,I,K). Our observation that A11.96 is the first and only A7.6-derived cell to co-express the siphon muscle determinants *Hand-r, Tbx1/10* and *Ebf* opened the possibility that A11.96 is the fate-restricted OSM precursor.

In order to determine whether the A11.96-derived A12.191/192 cells are the sole fate-restricted OSM founder cells, we characterized the expression dynamics of *Mrf* mRNA and the *Isl>unc76::Venus* reporter construct – hereafter referred to as *Isl>YFP*, which marks all siphon muscle cells (Fig. 1; (Stolfi et al. 2010)). Within the A7.6 lineage, we found that *Isl>YFP* was first expressed in A12.191/192 at 16hpf (Fig. 4A,A’). Subsequently, A12.192 and .191 cells each divide once between 18hpf and 22hpf, giving rise to four Isl>YFP+/LexO(A7.6)≫H2B:mCherry+ cells on either side of the larva (A13.382 to A13.386; Fig. 4A-D, with close-ups of the same data shown in A’-D’). These Isl>YFP+/LexO(A7.6)≫mCherry+ cells will migrate to form an 8-cell ring underneath the oral ectoderm (a.k.a. stomodeum; Christiaen et al, 2005), where at 24hpf the complete OSM rudiment will contain 8 cells, all of which are mCherry+/YFP+ (Fig.4D-D’).

Since *Mrf* and *Bhlh-tun1* were previously shown to be expressed in the 28hpf OSM ring (Razy-Krajka et al., 2014), we reasoned that the earliest expression of these genes would also point toward the OSMP. Therefore, we sought to describe their expression dynamics in the labeled A7.6 lineage. We found that *Mrf* first turned on at 18hpf in the most anterior pair of marked cells, which we inferred to be A12.191 and A12.192 (Fig. 4E). Between 18 and 22hpf, one of these cells divides first, so that at around 19hpf the OSMP are comprised of three cells—one *Mrf+/bHLH-tun1-;* one *Mrf-/Bhlh-tun1+;* and one expressing both genes (Fig. 4F). At later time points, we found that each set of OSMP divided into two *Mrf*-expressing and two *Bhlh-tun1-expressmg* cells (Fig. 4G, H). The existence of *Bhlhtunl+;Mrf-* OSMPs suggests that, by analogy with the ASMPs (Razy-Krajka et al. 2014), the OSM anläge contains stem-cell-like precursors that may contribute to siphon growth and/or regeneration later in life (e.g. Hamada et al., 2015). Taken together, these observations indicate that A7.6-derived fate-restricted OSM founder cells are A12.192 and A12.191, which express *Hand-r, Ebf, Tbx1/10, Isl>unc76::Venus* and *Mrf* before producing *Mrf+;Bhlh-tun1-*and *Mrf-;Bhlh-tun1+* OSMPs in a manner analogous to ASMPs (Fig. 4I, J).

### Novel regulatory relationships between *Tbx1/10* and *Ebf*

Because early determinants of ASM specification are expressed in the progenitors of the OSM, but follow different spatiotemporal dynamics, we sought to test whether *Tbx1/10* and *Ebf* interact functionally in specifying OSM and regulating each other's expressions. In the B7.5/cardiopharyngeal lineage, *Tbx1/10* function is required for *Ebf* expression, and contributes to inhibiting the heart program in the ASMs (Wang et al. 2013). However, because *Tbx1/10* is not observed in the OSMF until after *Ebf* has been expressed for several hours, we ruled out a role for *Tbx1/10* in the onset of *Ebf* expression but reasoned that *Tbx1/10* could instead contribute to *Ebf* maintenance in fate-restricted OSMPs.

We first tested whether *Tbx1/10* misexpression would maintain *Ebf* expression in A11.95-derived anterior TLCs that transiently express *Ebf* but fail to maintain it and never activate *Tbx1/10* (see Figs. 3L, 4I for the clonal context of this discussion). We used the LexO(A7.6)>>Tbx1/10 to overexpress Tbx1/10 throughout the A7.6 lineage, and analyzed the effect on *Ebf* expression at 16hpf (Fig. 5A-D). At 16hpf, *Ebf* is normally expressed in the nervous system, the ASMFs and the OSMPs (A12.192/191; Fig 31; Fig. 5B; Wang et al. 2013). Upon misexpression of *Tbx1/10*, we did not observe substantial ectopic *Ebf* expression in either the A11.95-derived anterior TLC, nor anywhere in the posterior TLCs (n=62; Fig. 5C). Thus Tbx1/10 misexpression does not appear to be sufficient to cause ectopic *Ebf* expression in the posterior TLC, or maintain *Ebf* expression in the anterior TLC. This is in contrast to the situation observed in the B7.5 lineage, where Tbx1/10 misexpression caused robust ectopic *Ebf* activation in the second heart precursors (Wang et al. 2013).

**Figure 5.**
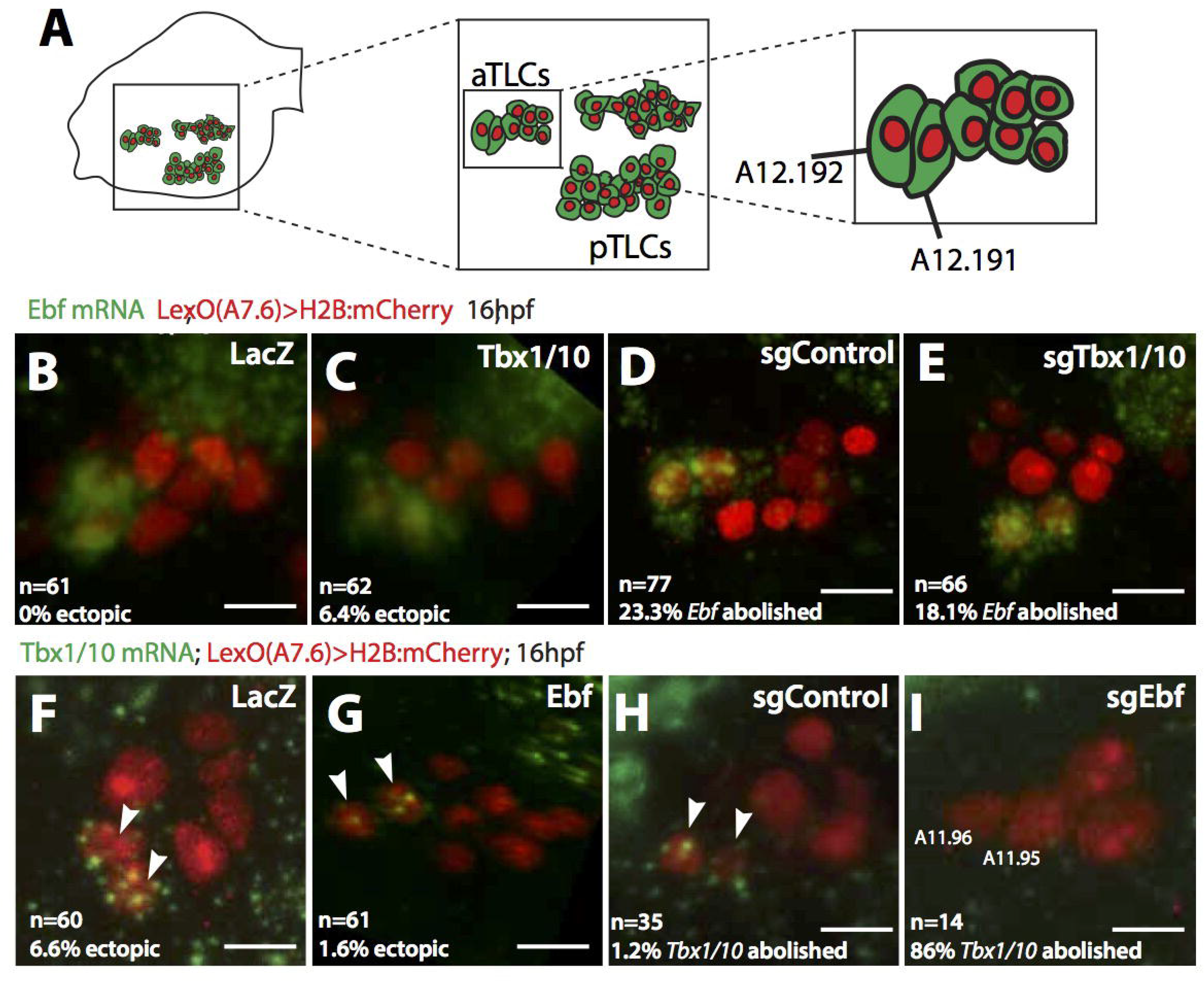
Effects of *Tbx1/10* gain-of-function and loss-of-function on expression of *Ebf (B-E)* and effect of *Ebf* gain-of-function and loss-of-function on *Tbx1/10* at 16hpf (F-I). (A) Cartoon showing whole-embryonic context of the anterior TLC at 16hpf. (B-E) Close-up of anterior TLC population at 16hpf, electroporated with LexO(A7.6)>H2B:mCherry and LexAop>LacZ (B); LexAop>Tbx1/10 (C); LexAop>nls:Cas9:nls;U6>sgControlF+E (D); or LexAop>nls:Cas9:nls;U6>sgTbx1.303;U6>sgTbx1.558 (E). (F-I) Close-up of anterior TLC in lóhpf embryos electroporated with LexO(A7.6)>H2B:mCherry and LexAop>LacZ (F); LexAop>Ebf (G); LexAop>nls:Cas9:nls;U6>sgControlF+E (H); or LexAop>nls:Cas9:nls;U6>sgTbx1.303;U6>sgTbx1.558 (I). Scale bars = 10μm. Total n are pooled from two biological replicates of batch-electroporation of zygotes. n=61 for (B); n=62 for (C); n=77 for (D); n=66 for (E).

We next sought to test the effects of loss of *Tbx1/10* on *Ebf* expression in the A7.6 lineage. In the B7.5 lineage, RNAi-mediated loss of Tbx1/10 function inhibited *Ebf* expression in the ASMF (Wang et al. 2013). To test whether Tbx1/10 is required for the maintenance *Ebf* expression in the OSMP, we used CRISPR/Cas9-based tissue-specific mutagenesis to inhibit *Tbx1/10* function. Here, we adapted methods developed by Gandhi, Stolfi, et al (Stolfi et al. 2014; Gandhi, Stolfi and Christiaen, in preparation) to direct editing of the *Tbx1/10* locus by CRISPR/Cas9 (see Materials and Methods and Fig. S2), and used a pair of sgRNA constructs targeting bp303-322 and bp558-577 in exon 1 of *Tbx1/10* in all *Tbx1/10* loss-of-function experiments.

Since RNAi experiments indicated that *Tbx1/10* is necessary for *Ebf* expression in the ASMF, we first tested the efficacy of CRISPR/Cas9-based genome editing of *Tbx1/10* in the B7.5 lineage by electroporating fertilized eggs with Mesp>nls:Cas9:nls and either U6>sgControl or U6>sgTbx1.303;U6>sgTbx1.558. Using *Ebf>GFP* as a proxy for *Ebf* activation (Wang et al. 2013), we found that *Tbx1/10* mutagenesis caused an 87% reduction in the proportion of electroporated larvae showing Ebf>GFP+ ASMs relative to control larvae, thus mimicking the published RNAi-mediated phenotype and demonstrating the efficacy *Tbx1/10*-targeting sgRNA constructs (Fig. S3B).

We then targeted Cas9 expression to the A7.6 lineage with our LexA/LexAop system and found that, in contrast to what has been demonstrated in the B7.5 lineage, loss of *Tbx1/10* function in the A7.6 lineage did not alter *Ebf* expression in the OSMPs at 16hpf (n=66; Fig. 5E). Taken together, these results indicate that, by contrast to its function in the B7.5 derived ASMF, *Tbx1/10* is not involved in either activating or maintaining *Ebf* expression in the A7.6-derived OSM precursors.

Next, we sought to test the role of *Ebf* in activation of *Tbx1/10* in the A7.6 lineage. It seems unlikely that *Ebf* alone activates *Tbx1/10* in the anterior TLC, since *Ebf is* expressed more broadly than *Tbx1/10*, and Ebf misexpression throughout the A7.6 lineage using a LexO(A7.6)>>Ebf strategy was not sufficient to cause ectopic *Tbx1/10* expression in any part of the TLC (Fig. 5F). However, *Ebf* may still be required together with (an) unknown co-factor(s) to activate *Tbx1/10* in the A11.96 OSM founder cells. To test this, we used published sgRNA constructs and LexO(A7.6)>>Cas9 for A7.6-lineage-specific CRISPR/Cas9-mediated targeted mutagenesis of the *Ebf* coding sequence (Stolfi et al. 2014). In control conditions (i.e. either LexO(A7.6)≫LacZ or LexO(A7.6)>>Cas9;U6>sgControl), *Tbx1/10* was expressed in the OSMP, sensory vesicle, STVC, and endoderm (Fig. S4). We found that Ebf-targeted mutagenesis inhibited *Tbx1/10* expression specifically in the OSMP at 16hpf in 86% of embryos (n=14; Figure 51). We therefore conclude that the OSM precursors uniquely require *Ebf* to express *Tbx1/10*, while *Ebf* expression is independent of *Tbx1/10* expression. This is in stark contrast with the regulatory relationship between *Tbx1/10* and *Ebf* in the ASM, where *Tbx1/10* is required for *Ebf* expression and misexpression of *Tbx1/10* was sufficient to cause ectopic *Ebf* expression within the cardiopharyngeal mesoderm (Wang et al. 2013).

### Tbx1/10 and Ebf are required in parallel to specify the OSM fate

We have shown that, although ASM and OSM both express *Mrf* in the siphon muscle founder cells, the upstream regulators of *Mrf* expression in B7.5-derived ASMP are deployed differently in the A7.6 lineage. The regulatory relationships that we have demonstrated between *Ebf* and *Tbx1/10* open the possibility that the function of each in *Mrf* activation may also be unique in OSM vs. ASM. We first tested whether loss of either *Ebf* or *Tbx1/10* function throughout the A7.6 lineage inhibited OSM fate specification by assaying *Mrf* expression in 24hpf larvae (Fig. 6). In samples electroporated with

**Figure 6.**
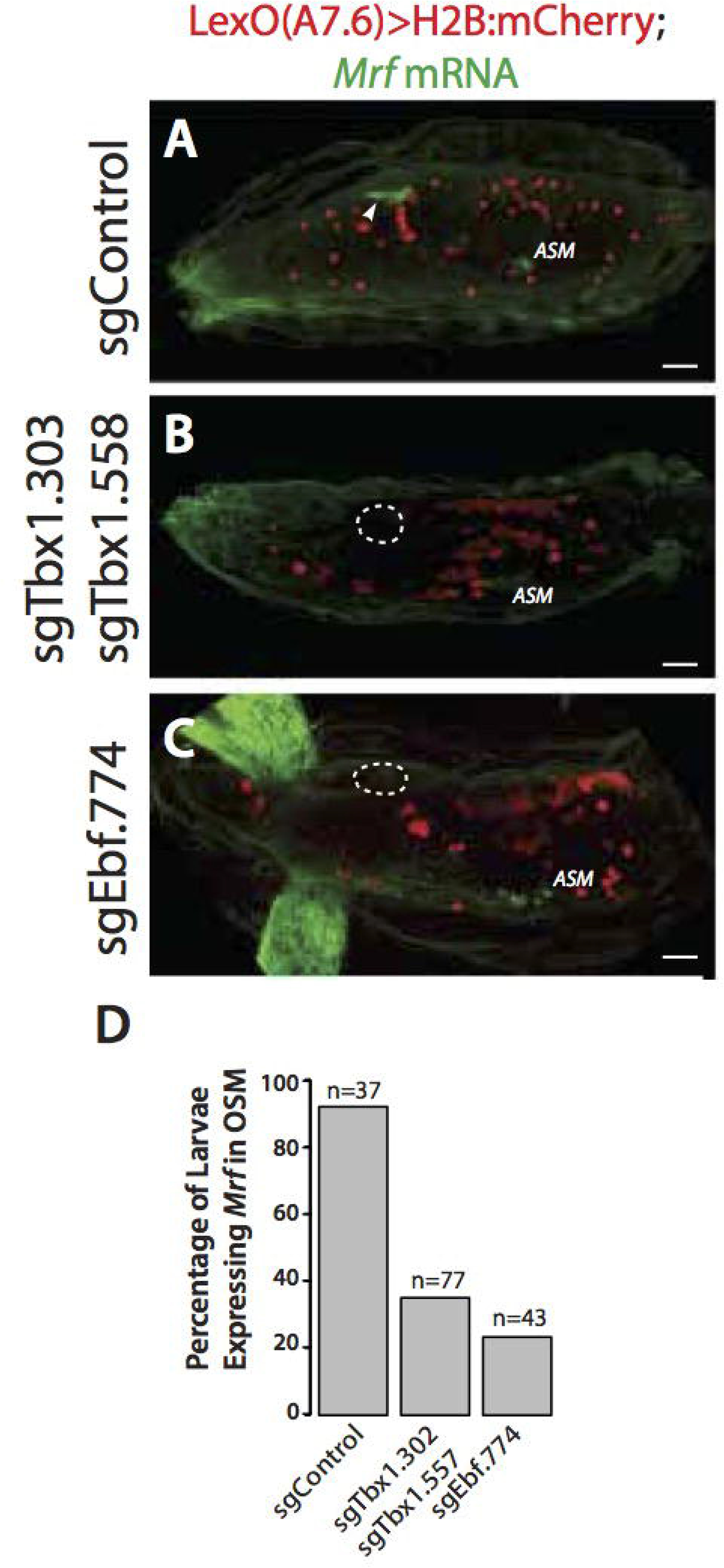
Tissue-specific knockdown of *Ebf* or *Tbx1/10* in the A7.6 lineage leads to loss of OSM. (A-C) 26hpf larvae electroporated with LexO(A7.6)>H2B:mCherry;LexAop>nls>Cas9:nls and U6>sgControlF+E (A), U6>sgTbx1.303;U6>sgTbx1.558 (B), or U6>sgEbf.774 (C). (D) Boxplot showing the proportion of larvae in which LexO(A7.6)>H2B:mCherry and *Mrf* mRNA were both expressed in the OSM, with sample sizes indicated. The total n are pooled from two biological replicates of batch-electroporation of zygotes: n=37 for sgControl; n=77 for sgEbf; n=43 for sgTbx1/10. Scale bars = 25μm.

LexO(A7.6)≫nls:Cas9:nls and control sgRNA (U6>sgControlF+E; Stolfi et al, 2014), *Mrf* was co-expressed with LexO(A7.6)>>H2B:mCherry in the OSM of 91.9% of larvae (n=37; Fig. 6A, D). Co-electroporating the sgTbx1.303 and sgTbx1.558 constructs targeting *Tbx1/10* reduced the proportion of transfected larvae expressing *Mrf* in the vicinity of the oral ectoderm to 35.1% (n=77; Fig. 6B, D). Similarly, targeted mutagenesis of *Ebf reduced* the proportion of larvae with *Mrf* +/LexO(A7.6)>>mCherry+ OSM to 23.3% (n=43; Fig. 6C, D). Because *Tbx1/10* does not appear to be involved in *Ebf* activation (Fig. 5), we interpret these results to indicate that *Tbx1/10* has a direct, *Ebf*-independent role upstream of *Mrf* in OSM fate. On the other hand, since *Ebf is* necessary for *Tbx1/10* expression in the OSM (Fig. 5), loss of *Mrf* upon *Ebf* mutagenesis could be due to loss of *Tbx1/10* in *cis*. Therefore, these data indicate that although the role of *Ebf* and *Tbx1/10* as siphon muscle regulators is conserved between OSM and ASM, these functions may result from different regulatory logic in the two contexts.

We sought to further test this possibility using gain-of-function assays by misexpression. We used *LexO(A7.6)≫Ebf* or *LexO(A7.6)≫Tbx1/10* to over-express Ebf and/or Tbx1/10 throughout the A7.6 lineage and assayed *Mrf* expression in 24hpf larvae (Fig. 7). Using LexO(A7.6)≫LacZ as a control, we observed that 4.9% of larvae express low levels of *Mrf* among scattered mesenchymal cells (n=41; Fig. 7A, E). Similarly, misexpression of *Tbx1/10* resulted in only 2.1% of larvae showing any ectopic *Mrf* expression (n=47; Fig. 7B, E). Meanwhile, misexpression of *Ebf* increased the proportion of larvae with ectopic *Mrf* to 32.8% (n=64; Fig. 7C, E). In these larvae, A7.6-derived cells that showed ectopic *Mrf* expression clustered towards the dorsal midline. We observed increased proportions of ectopic *Mrf+* A7.6-derived cells when we co-expressed LexO(A7.6)≫Ebf and LexO(A7.6)≫Tbx1/10 together: 60.7% of larvae showed ectopic *Mrf* expression (n=84; Fig. 7D, E). Remarkably, A7.6-lineage cells that ectopically expressed *Mrf* upon *Ebf* and *Tbx1/10* co-expression tended to cluster near the atrial siphon primordium, whereas the anterior-born OSMP normally migrate towards the oral siphon primordium (a.k.a. stomodeum; e.g. Fig. 7D). This observation suggests that when combined misexpression of *Ebf* and *Tbx1/10* induced ectopic *Mrf* expression and siphon muscle fate in the posterior TLCs, siphon muscle precursor cells home towards the closest siphon primordium during larval development. Taken together, these data demonstrate that *Ebf* and *Tbx1/10* functions are both required and act in parallel to activate *Mrf* and promote siphon muscle specification in the A7.6 lineage.

**Figure 7.**
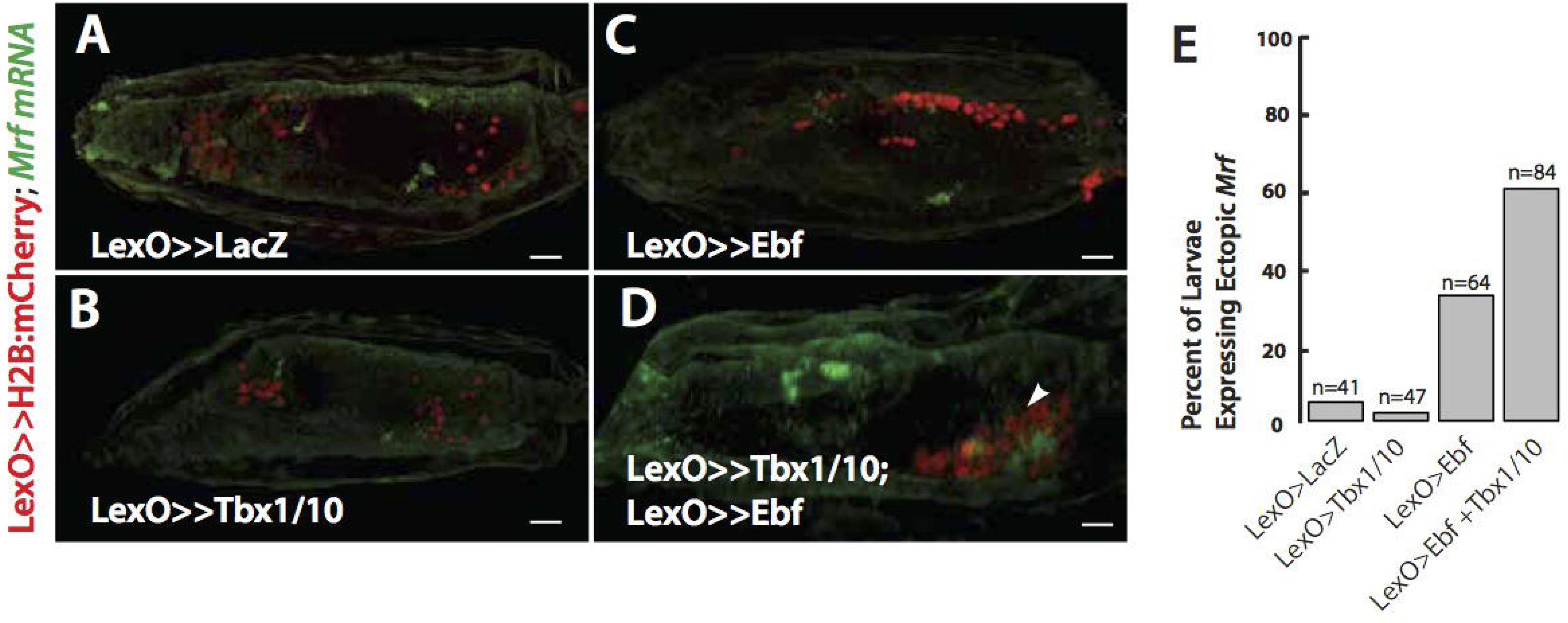
Effect of *Ebf* or *Tbx1/10* gain-of-function on expression of *Mrf* (A-D) 24hpf larvae electroporated with LexO(A7.6)>>H2B:mCherry and LexAop>LacZ (A); LexAop>Ebf (B); LexAop>Tbx1/10 (C); or LexAop>Ebf; LexAop>Tbx1/10 (D). (E) Boxplot showing proportion of larvae in each condition in which we observed ectopic *Mrf* expression, with sample sizes indicated. n= 41 for LacZ; n=47 for Ebf; n=64 for Tbx1/10; n=84 for Ebf + Tbx1/10.

### Context-specific wiring of a conserved siphon muscle differentiation kernel

Having established the parallel requirement for *Ebf* and *Tbx1/10* upstream of *Mrf* in OSM specification, we sought to test whether this regulatory architecture also governs ASM specification. Previous work suggested that *Ebf is* necessary and sufficient for ASM specification in the B7.5 lineage, where it must be activated by *Tbx1/10*. In order to determine whether *Ebf is* sufficient for ASM fate specification in the absence of *Tbx1/10*, we designed a strategy using CRISPR/Cas9 to mutate *Tbx1/10* in the B7.5 lineage; while using a minimal B7.5-lineage-specific *Tbx1/10* enhancer (Racioppi et al., unpublished construct) to restore *Ebf* expression specifically in the STVCs and their progeny. Importantly, the enhancer activity of this Tbx1/10 construct was not affected by expression of Cas9 and sgTbx1.303;558 (Fig. S3A, B; *Tbx1/10(-7333/-2896)::bpTbx1>Ebf)*.

In control larvae, there are four ASM precursors (ASMP) at 22hpf, with the outer ASMP expressing *Mrf* and the inner ASMP expressing *Bhlh-tun1* ((Razy-Krajka et al. 2014) and Fig. 8A; SHP marked with “*”). In the B7.5 lineage, loss of *Tbx1/10* results in loss of *Ebf* in the ASMF (Fig. S3C; Wang et al., 2013) and Ebf activates ASMP-specific expression of Mrf and Bhlh-tun1 (Razy-Krajka et al., 2014). Consistent with this, we found that loss of *Tbx1/10* abolished expression of *Mrf* and *Bhlh-tun1* in 100% of larvae (n=10), likely due to loss of *Ebf* expression (Fig. 8C). Because the *Tbx1/10* enhancer drives transgenic expression in the ASM founder cells (ASMF) as well as the second heart precursors (SHP; see Fig. 4J), we expected to see the effects of *Ebf* overexpression expanded to the SHP. Indeed, electroporation of *Tbx1/10>Ebf* with control guide RNA caused an expansion of *Mrf* and *Bhlh-tun1* expression to the SHP in 9 out of 11 larvae (Fig. 8B). Having also confirmed that mutation of *Tbx1/10* did not interfere with expression of *Tbx1>GFP* (Fig. S3B), we introduced *Tbx1>Ebf* and found that *Ebf* alone was able rescue, and cause ectopic, expression of *Mrf* and *Bhlh-tun1* (Fig. 8D). These data indicate *Tbx1/10* function is dispensable for ASM specification downstream of *Ebf*. Therefore, whereas *Ebf can* regulate siphon muscle fate without *Tbx1/10* in the ASM, its function as an activator of siphon muscle fate in the OSM depends on co-expression with *Tbx1/10* (Fig. 8E).

**Figure 8.**
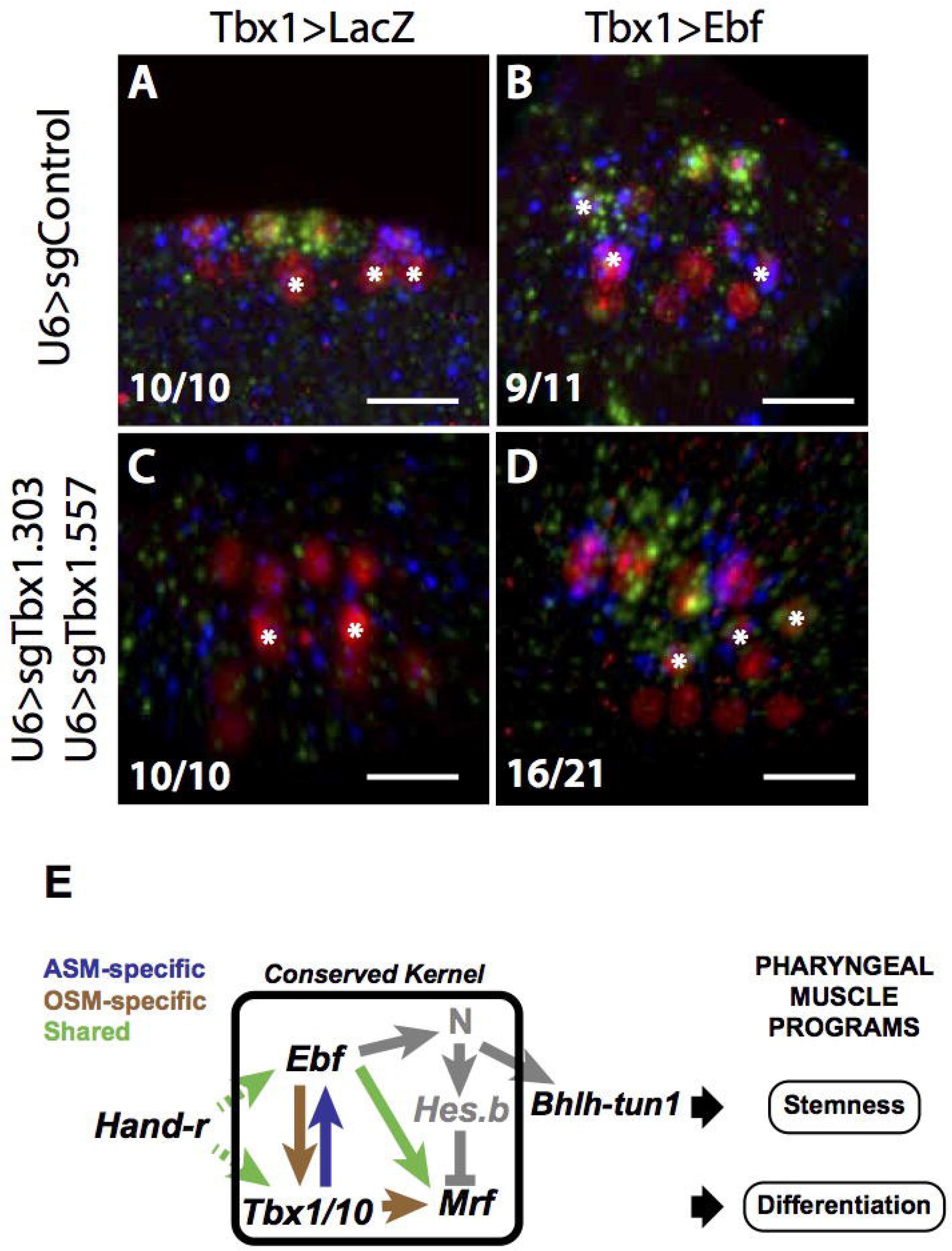
*Ebf is* an independent master regulator of siphon muscle fate in the B7.5 lineage. (A-D) Close-up of ASM and first and second heart precursors (SHP; marked with “*”), derived from the B7.5 lineage, revealing *Mrf* mRNA (blue) and *Bhlh-tun-1* mRNA (green). (A,B) Larvae electroporated with Mesp>H2B:mCherry;Mesp>nls:Cas9:nls;U6>sgControlF+E and Tbx1>LacZ (A) or Tbx1>Ebf (B). In (B), note that expression of *Mrf* and *Bhlh-tun-1* has expanded to the SHP, due to earlier Tbx1>Ebf expression in the secondary TVCs. (C, D) Larvae electroporated with Mesp>H2B:mCherry;Mesp>nls:Cas9:nls;U6>sgTbx1.303;U6>sgTbx1.558 and Tbx1>LacZ (C) or Tbx1>Ebf (D). Note that in (C) there is a complete loss of *Mrf or Bhlh-tun-1* expression, whereas in (D), *Mrf* and *Bhlh-tun-1* show wild-type expression patterns in the ASM, and are also expressed in the SHP. Scale bars = 25μM. n=. All data was collected from a single technical replicate. (E) Schematic diagram comparing core regulatory interactions upstream of *Mrf*-driven differentiation and *Notch*-driven stemness in ASM and OSM. In black, documented shared expression of *Hand-r* in cells that give rise to, among other tissues, siphon muscles may be involved in activation of the core common regulators *Tbx1/10* and *Ebf*, as indicated by green dashed arrows. Although both *Ebf* and *Tbx1/10* impinge on *Mrf* expression, the distinct regulatory relationships in place in ASM vs. OSM are indicated by purple and brown arrows, respectively. The shared direct input from *Ebf to Mrf in* both ASM and OSM is indicated by the solid green arrow. In grey, Notch signaling downstream of *Mrf* activation has been established in the ASM (Razy-Krajka *et al*. 2014) as the mechanism for cells to choose between stemness and differentiation, but has not yet been tested in the OSM.

## Discussion

In this paper, we use two clonally distinct but molecularly similar muscle groups to examine context-dependent control of muscle specification. Though the regulatory mechanisms upstream of *Mrf* expression in the atrial siphon muscle (ASM) founder cells of the basal Chordate *Ciona intestinalis* have been described (Stolfi et al. 2010; Wang et al. 2013; Razy-Krajka et al. 2014), very little was known about the origins of the oral siphon muscles (OSM) or the mechanisms of OSM specification. We present a detailed description of the developmental origins of the oral siphon muscles in the ascidian *Ciona*. We have traced the clonal origins of the OSM from a single multipotent mesodermal progenitor in the gastrula, and explored the mechanisms that activate myogenesis in this lineage. In order to identify and characterize the descendants of the A7.6 lineage, we adapted the heterologous Gal80-repressible binary transgenic system from *E. coli*, LexA/LexAop (R. Yagi, Mayer, and Basler 2010), which is poised to permit refined transgenic strategies for studies using *Ciona*.

### Conserved synergy between *Ebf* and *Tbx1/10*

Muscle differentiation across bilaterians depends on the activity of muscle regulatory factor (MRF) family of bHLH DNA binding transcription factors (Braun et al. 1994; Summerbell, Halai, and Rigby 2002; Kassar-Duchossoy et al. 2004; Rudnicki et al. 1993; Wei, Rong, and Paterson 2007; Fukushige and Krause 2005; Andrikou et al. 2015). These MRFs are regulated by, and work in concert with, transcription factors that can differ between muscle groups, but appear to be conserved across species. Pax3/7 homologs have long been recognized as regulators of somitic muscle specification and regeneration in vertebrates, and have recently been shown to have a role in limb muscle specification and regeneration in arthropods (Buckingham and Relaix 2007; Sambasivan et al. 2013; von Maltzahn et al. 2013; Konstantinides and Averof 2014). *Tbx1* homologs are crucial regulators of pharyngeal muscle development in vertebrates (Aggarwal et al. 2010; Z. Zhang, Huynh, and Baldini 2006; Sambasivan et al. 2009; Diogo et al. 2015). In mammals, *Tbx1* regulates *Myf5* and *MyoD* specifically in the mandibular arches, but is not involved in specification of other head muscles, or in the somitic muscles (Sambasivan et al. 2009). The *Drosophila* ortholog of Tbx1, org-1, is required for expression of *ladybird* and *slouch*, both muscle identity genes, all of which occurs downstream of *nautilus* expression (Crozatier and Vincent 1999; Wei, Rong, and Paterson 2007; Schaub et al. 2012).

Transcription factors of the Collier/Olf1/Ebf (COE) family are emerging as important upstream regulators of *Mrf* expression and myogenesis. *COE* orthologs have documented roles in neurogenesis, cellular immunity, and hematopoiesis (Crozatier and Vincent 1999; Pang, Matus, and Martindale 2004; Kratsios et al. 2011; Benmimoun et al. 2015). Earliest evidence for *COE’s* myogenic properties implicated it in specification of a hemisegmentally-repeated abdominal muscle subtype by interactions with nau,though in a domain distinct from the action of *org-1* (Crozatier and Vincent 1999; Schaub et al. 2012; Wei, Rong, and Paterson 2007) (Crozatier and Vincent 1999). The COE homologs, Ebf2 and −3, have been found to regulate *Myf5* and *MyoD* expressions in *Xenopus* (Green and Vetter 2011). In the mouse, Ebf1 and Ebf3 interact with MyoD in the developing diaphragm to activate muscle-specific gene transcription (Jin et al. 2014). *Ciona* has only one copy of *Ebf*, which has been found to be necessary for both neurogenesis and myogenesis (Kratsios et al. 2011; Razy-Krajka et al. 2014), in keeping with an ancient role for *COE* homologs in both of those functions (Jackson et al. 2010).

Future studies in other systems will be needed to determine whether, as shown to be the case in *Ciona* (this study and Wang et al. 2013; Razy-Krajka et al. 2014), regulatory interactions between COE and Tbx1 contribute to muscle specification as could be predicted by their overlapping expressions in developing pharyngeal muscles.

### Comparison of ASM and OSM

We have shown that, while *Tbx1/10* and *Ebf are* both expressed in the ASM and OSM precursors and required for *Mrf* expression, their regulatory relationships differ between the B7.5/ASM and A7.6/0SM lineages (Fig. 8). In the B7.5 lineage, *Ebf is* necessary and sufficient to activate *Mrf even* in the absence of *Tbx1/10* (Fig. 8; Wang et al. 2013; Razy-Krajka et al. 2014). In the A7.6 lineage, on the other hand, *Mrf* requires the combined inputs of *Ebf* and *Tbx1/10* (Fig. 6 and 7). When misexpression of *Ebf* leads to ectopic *Mrf* expression (Fig. 7), it was likely due to the ability of *Ebf* to activate *Tbx1/10* in the A7.6 lineage (Fig. 5), so that *Tbx1/10* was provided in *trans*, and both together activated *Mrf*. Ectopic activation of *Mrf in* the A7.6 lineage was nearly twice as efficient when *Tbx1/10* and *Ebf were* provided together, supporting the notion that they act in combination to activate *Mrf in* the OSM.

Why would the temporal deployment and regulatory logic of two such deeply conserved muscle-regulatory genes differ so greatly in each context? Specification of the siphon muscle founder cells in both the A7.6 lineage and the B7.5 lineage is an instance of binary fate choice, the outcome of which is expression of *Mrf* and the terminal differentiation gene battery specific to siphon muscles. However, the alternative fate and embryonic context in each case is different. Therefore, the rewiring of the *Ebf-Tbx1/10* interactions that we have demonstrated upstream of *Mrf* may reflect larger network constraints specific to the A7.6 and B7.5 lineage, respectively. Meanwhile, the shared need for either *Tbx1/10* or *Ebf in* muscle specification likely reflects the fact that *Ebf* and *Tbx1/10* regulate not only *Mrf* but is also involved in regulating terminal differentiation genes in concert with *Mrf*. Such cooperation between COE and MyoD to activate muscle-specific differentiation genes has already been demonstrated in mouse (Jin et al. 2014). *COE* in *Drosophila* has also been shown to bind directly to enhancers of muscle-specific identity genes (de Taffin et al. 2015), as has *org-1* (Schaub et al. 2012). Our results indicate that although *Ebf* has been largely overlooked as a muscle regulator in vertebrates (perhaps because of genetic redundancies between Ebf1, −2 and −3), its function in concert with MyoD homologues may be deeply conserved, and medically relevant. Further work investigating genes directly activated by *Ebf in* ASM and OSM in *Ciona*, will provide important insights into the conservation and evolution of the complex regulatory interactions involved in muscle development.

## Materials and Methods

### Animals and Electroporation

Gravid *C. intestinalis* type A, also known as *Ciona robusta* (Brunetti et al. 2015) adults were obtained from M-REP, Santa Barbara, CA. Collection of gametes, fertilization, dechorionation, and electroporation of zygotes were all performed as described previously (Christiaen et al. 2009b). Animals were electroporated with 10-60 *μ* g of plasmid DNA, and raised at 18°C.

### Cloning of unary transgenic enhancers

Enhancers were amplified from larval gDNA using specific primers shown in Table 1 and cloned into backbones containing reporters using standard molecular cloning procedures.

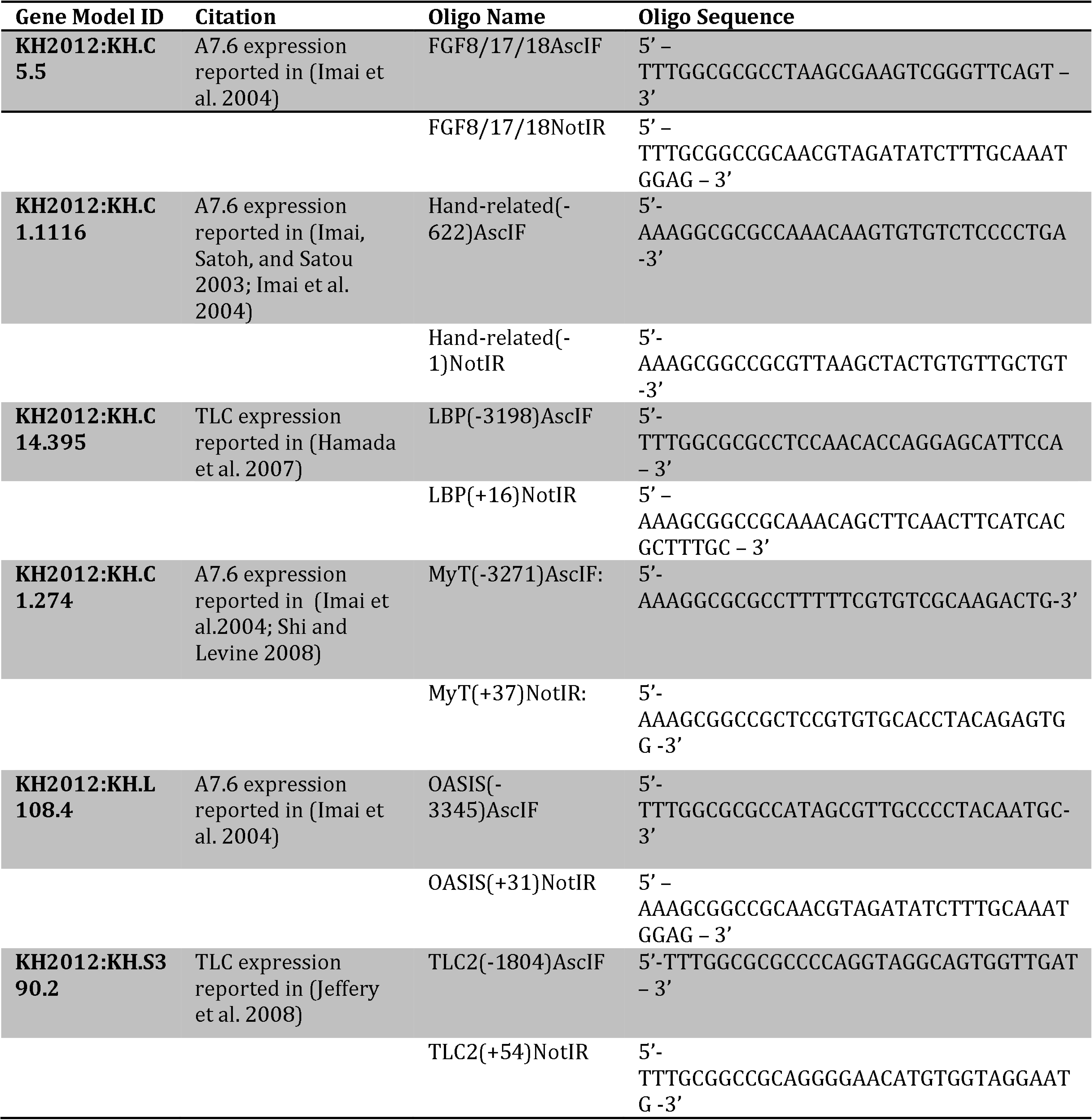

### Cloning of Gal4/UAS, LexA/LexAop, TrpR/tUAS and QF/QS constructs

Gal4 was amplified from Tubp-Gal4 (Addgene #17747; (Lee and Luo 1999)) using specific primers and adding NotI and EcoRI restriction sites on the 3’ and 5’ ends, respectively (NotI-Gal4-F: 3’- ATGAAGCTACTGTCTTCTATC −5’; EcoRI-Gal4-R: 3’-TTACTCTTTTTTTGGGTTTGG −5’). Gal80 was amplified from Tubp-Gal80 (Addgene #17748; (Lee and Luo 1999)) using specific primers and adding NotI and EcoRI restriction sites (NotI-Gal80-F 5’- AAAGCGGCCGCAACCATGGACTACAACAAGAGATC −3’; EcoRI-Gal80-R 5’- AAAGAATTCTTATAAACTATAATGCGAGAT −3’). Plasmids containing a 5x concatamer of the UAS response element fused to the HSP70 basal enhancer were created by excising CD8:GFP from the pUASt-mCD8:GFP backbone (Addgene #17746; (Lee and Luo 1999)) using NotI/XbaI restriction digest, and then using an XbaI/EcoRI linker fragment to clone all reporters flanked by NotI/EcoRI sites.

The LHG coding sequence and LexAop enhancer sequences were taken from *pDPPattB-LHG* and *p28-pJFRCl 9-13xLexAop2-IVS-myr-GFP* (R. Yagi, Mayer, and Basler 2010). The LHG coding sequence was amplified using specific primers and adding restriction sites NotI/EcoRI (NotI-LHG-F: 5’- AAAGCGGCCGCAACCATGAAAGCGTTAACGGCCAG −3’; EcoRI-LHG-R 5’-TTTGAATTCTTACTCTTTTTTTGGGTTTGGT −3’ and cloned downstream of the Hand-r(-622/-1) enhancer. The 13xLexAop enhancer along with the *Drosophila melanogaster* HSP70 basal promoter was excised from the *pJFRC19-13xLexAop2-IVS-myr-GFP* vector using AscI/NotI restriction enzymes and cloned into a backbone containing H2B::mCherry and subsequently subcloned using standard molecular cloning methods.

TrpR coding sequence was amplified from pCMV:nlsTrpR-Gal4AD using specific primers and adding NotI and EcoRI restriction sites at the 3’ and 5’ ends, respectively (NotI-TrpR-F: 3’- AAAGCGGCCGCAACCATGGCACCCAAGAAGAAGAGGAAG – 5’; EcoRI-TrpR-R: 3’-GCCCAACAATCACCCTATTCAGC – 5’)

The QF coding sequence was amplified from pAC-QF (Addgene #24338; (Potter et al. 2010)) using specific primers and adding NotI and EcoRI restriction sites at the 3’ and 5’ ends, respectively (NotI-QF-F: 3’-AAAGCGGCCGCAACC ATGCCGCCTAAACGCAAGAC; EcoRI-QF-R: 3’-AAAGAATTC CTATTGCTCATACGTGTTGAT-5’). The QUAS response element and *D. melanogaster* HSP70 basal promoter was amplified from p5E-QUAS (Addgene #61374) using specific primers and adding NotI and EcoRI restriction sites at the 3’ and 5’ ends, respectively (NotI-QUAS-F: 3’ – TTT GGCGCGCC GGG TAA TCG CTT ATC CTC GG – 5’; EcoRI-QUAS-R: 3’- TTT GCGGCCGC CAA TTC CCT ATT CAG AGT TCT – 5’).

### Genome editing using CRISPR

Single Guide RNAs against *Ebf were* used according to the methods described in (Stolfi et al. 2014). Single guide RNAs against *Tbx1/10 were* designed using CRISPRdirect (Naito et al. 2014), but eliminating putative targets that fell on SNPs documented in the published genome. Complementary oligonucleotides of each (N)21-GG target were synthesized by Sigma-Aldrich, St. Louis, Missouri, USA, and cloned downstream of the U6 RNA polymerase III promoter according to the methods described in (Stolfi et al. 2014). We tested a total of nine putative sgRNAs (“+” or “-” indicates whether the PAM was on the + or – strand; the number after the period indicates the nucleotide number on the cds of the first base-pair targeted): sgTbx1.303(+); sgTbx1.422(-); sgTbx1.558(+); sgTbx1.673(+); sgTbx1.783(+); sgTbx1.835(+); sgTbx1.982(+); sgTbx1.971(-); sgTbx1.1067(+) (Fig. S2A). We verified that sgRNAs directed cutting of genomic DNA by electroporating embryos with the ubiquitously-expressed Ef1 *α* >nls:Cas9:nls and 25 *μ* g each of a single U6>sgTbx1 plasmid. We extracted gDNA from 17hpf larvae, amplified the targeted region using specific primers, and TOPO-cloned the PCR products into pCR-II vectors for sequencing. We found that when electroporated individually, sgTbx1.303 and sgTbx1.558 produced the most efficient cutting, with 2/6 and 4/8 sequenced clones, respectively, showing mutations in the targeted region. However, when paired, sgTbx1.303 and sgTbx1.558 produced a large deletion in 6/8 clones sequenced (Fig. S2B). Therefore, for all tissue-specific manipulations of Tbx1/10 function, we used LexO(A7.6)≫nls:cas9:nls;U6>sgTbx1.303;U6>sgTbx1.558, with 25ug of Hand-r>LHG, 25ug of LexAop>nls:Cas9:nls, and 25ug each of U6>sgTbx1.303 and U6>sgTbx1.558, referred to as LexO(A7.6)>sgTbx1 throughout the text.

## Acknowledgements

We are grateful to Anthony Filipovic for help characterizing the efficacy and toxicity of various candidate binary systems. We are indebted to Claudia Racioppi and Alberto Stolfi for the *Tbx1/10* STVC-specific enhancer. We give special thanks to Maximilien Courgeon and Claude Desplan for the LexA/LexAop plasmids. This work was supported by National Institutes of Health/National Heart, Lung and Blood Institutes R01 HL108643 award to L.C. and 2T32HD007520-16 award to TRT as part of the New York University Developmental Genetics Training Grant program (P.I.: Jessica Treisman). The authors declare no competing interests.

